# A connectome-based prediction model of long-term memory

**DOI:** 10.1101/2020.09.02.279372

**Authors:** Qi Lin, Kwangsun Yoo, Xilin Shen, R Todd Constable, Marvin M Chun

**Affiliations:** Department of Psychology, Yale University; Department of Radiology and Biomedical Imaging, Yale School of Medicine; Interdepartmental Neuroscience Program, Yale University; Department of Neurosurgery, Yale School of Medicine; Department of Neuroscience, Yale School of Medicine

## Abstract

Although many studies have investigated the neural basis of intra-individual fluctuations in long-term memory (LTM), few have explored the differences across individuals. Here, we characterize a whole-brain functional connectivity (FC) network based on fMRI data in an n-back task that robustly predicts individual differences in LTM. Critically, although FC during the n-back task also predicted working memory (WM) performance and the two networks had some shared components, they are also largely distinct from each other: the LTM model performance remained robust when we controlled for WM and vice versa. Furthermore, regions important for LTM such as the medial temporal lobe did contribute, but only partially, to predicting LTM. These results demonstrate that individual differences in LTM are dependent on the configuration of a whole-brain functional network including but not limited to regions traditionally associated with LTM during encoding and that such a network is separable from what supports WM.

## 1 Introduction

The ability to hold information in long-term memory (LTM) varies dramatically across individuals: while some can instantaneously recognize a stranger whom they talked to at a party 20 years ago, some struggle to even remember a classmate they met earlier that week. Why is this the case? Although there have been a rich body of work on the neural mechanism underlying within-subject fluctuations in LTM performance, such as the subsequent memory effect (Keerativittayayut et al., 2018; Kim, 2011; Paller & Wagner, 2002; Wagner et al., 1998), to date, only a few studies have investigated the neural basis of individual differences in LTM performance across subjects (Jangraw et al., 2018), most of which focused on the aging population (Meskaldji et al., 2016; Trelle et al., 2020). However, even among young adult population, there remains a large variance in LTM performance (for a recent review, see Unsworth, 2019). The neural basis for such variability remains unclear.

In stark contrast to the relatively sparse literature on the between-subject differences in LTM, there has been a burgeoning body of work characterizing the neural correlates of individual differences in working memory (WM), the ability to maintain and manipulate a limited amount of information actively over a short period of time (for a review, see Luck and Vogel, 2013). In fMRI studies of WM, the activity in posterior intraparietal sulcus (IPS) scales with WM load until the individual subject’s WM capacity is reached (Todd & Marois, 2004, 2005; Xu & Chun, 2006) and the difference in IPS activation between high and low set sizes is correlated with WM capacity (Todd & Marois, 2005). In addition to the activation difference in IPS, preparatory activities in the prefrontal cortex and basal ganglia were also associated with individual differences in WM capacity (McNab & Klingberg, 2008). More recent work focuses on functional connectivity, a measure of how synchronized different brain regions are, and characterizes whole-brain networks that are predictive of WM capacity within young adult populations and generalizable to older people or psychiatric populations (Avery et al., 2019; Bertolero et al., 2018; Yamashita et al., 2018). The intense interest in individual differences in WM and its neural basis perhaps comes as no surprise given the foundational role of WM in supporting a suite of more complex cognitive abilities (such as reasoning, fluid intelligence and planning; Conway et al., 2003; Süß et al., 2002) and its impairment in a range of psychiatric populations (such as schizophrenia, autism and major depression: Forbes et al., 2008; Lever et al., 2015; Snyder, 2013). Considering that a moderate correlation is also often observed behaviorally between WM and LTM (Unsworth, 2010) and that WM is central to such a wide range of cognitive functions, could the individual variability in LTM be explained by the neural network underlying individual differences in WM?

The similarity and distinction between memory on different timescales is probably one of the oldest questions in memory research (Cowan, 2008; James, 1891). Despite the phenomenological differences of these two kinds of memory, there remain controversies about whether they are supported by distinct neural mechanisms, mostly centering around the medial temporal lobe (including the hippocampus) and whether these regions are necessary for both WM and LTM. Early studies about patients such as H.M. show that damage to the medial temporal lobe leaves the patients’ WM intact but impairs their ability to form new long-term memory, suggesting the medial temporal lobe play a critical role in LTM but not WM (e.g., Scoville and Milner, 1957). However, more recent work found activation in the medial temporal lobe during WM task (Ranganath & D’Esposito, 2001). In addition, active maintenance of items during the early stage of WM delay contributes to later LTM formation, modulated by a network including dorsolateral PFC and the hippocampus (Blumenfeld & Ranganath, 2006; Ranganath et al., 2005). In line with these results in healthy populations, patients with medial temporal damage showed impairments in WM for visual stimuli (Olson et al., 2006a) and conjunctions (Olson et al., 2006b). These results have led some researchers to cast doubt on the double dissociation between WM and LTM (for reviews, see Graham et al., 2010; Jeneson and Squire, 2012; Ranganath and Blumenfeld, 2005).

Although these previous works have made valuable contributions to understanding the neural mechanisms behind WM and LTM, they tend to focus on group level analysis and a circumscribed set of brain regions. To fully understand the functional architectures supporting these two memory functions, we need to consider the factors affecting the variation of WM and LTM across individuals (Dubois & Adolphs, 2016; Gabrieli et al., 2015). Theoretically, such findings will complement past studies on group level focusing on the mechanism consistent across individuals and may reveal novel neural factors critical for these functions. Clinically, focusing on individual differences can provide insights for developing neuromarkers of memory deficits observed in psychiatric or aging populations.

Recent developments in connectome-based predictive modeling (CPM; Shen et al., 2017) have validated the utility of building predictive models of different cognitive functions and traits such as fluid intelligence, sustained attention, extraversion and working memory from whole-brain functional connectivity (e.g., Avery et al., 2019; Finn et al., 2015; Hsu et al., 2018; Rosenberg et al., 2015). A distinct advantage of the CPM framework lies in its flexibility: based on the same functional connectivity data, we can build different models that are predictive of a range of behaviors. By comparing these models, we can potentially identify the dissociable brain networks that support separable functions (e.g., IQ and sustained attention: Rosenberg et al., 2015) or overlapping brain networks underlying functions that may be related to each other (e.g., IQ and WM: Avery et al., 2019; sustained attention and reading: Jangraw et al., 2018). This flexibility of CPM has made it an ideal approach to probe the similarity and distinction between WM and LTM networks.

In the current study, we leverage the CPM framework to investigate the neural basis of individual differences in LTM and ask whether WM and LTM share the same neural substrates or whether they are largely distinct from each other. Additionally, we examine whether brain regions associated with intra-individual fluctuations in LTM also underlie between-individual differences. Using CPM to analyze data from the Human Connectome Project (Essen et al., 2013), we first characterized a predictive model of individual differences in LTM based on whole-brain functional connectivity during a WM task (n-back). We then compared the LTM network with a separate WM network built to predict 2-back accuracy in the same group of subjects based on the same whole-brain FC data (see also Avery et al., 2019). Furthermore, we explored the utility of regions previously associated with LTM formation such as the medial temporal lobe and regions showing subsequent memory effects in predicting individual differences in LTM.

## 2 Results

To investigate the neural basis of individual differences in long-term memory, we used data from the S1200 release of the Human Connectome Project (HCP; Essen et al., 2013; N = 708, see Methods for inclusion criteria). Specifically, we focused our analyses on the two runs of 5-minute n-back tasks during fMRI^1^ and the corresponding LTM and WM measures. LTM was measured using *d’* in a recognition memory task with the images presented during the n-back fMRI runs. The recognition memory task was performed outside of the scanner after all the fMRI runs during the session with the n-Back tasks. WM was measured using 2-back accuracy during the n-back tasks.

To explore whether FC during n-back tasks predicts memory performance, we conducted analysis following the Connectome-based Predictive Modeling pipeline (CPM; Finn et al., 2015; Rosenberg et al., 2015; Shen et al., 2017). For each n-back run, fMRI data were parcellated into 368 nodes (Shen et al., 2013) and a mean time course was calculated for each node. A whole-brain FC matrix was then created by calculating the pairwise Pearson’s *r* correlation and transforming these correlations via Fisher’s *z*, yielding a 368*×*368 matrix per each run for each participant. The two matrices calculated from the two n-back runs were then averaged to produce one matrix per participant.

These matrices were then submitted to a ridge-regression-based version of CPM (Gao et al., 2019; Greene et al., 2020) to predict LTM and WM separately. In brief, using 10-fold cross-validation, we first identified edges that were most correlated (*p* < 0.01) with the target behavior (LTM or WM) in the training sample and built a ridge regression model to predict the target behavior using these selected features. The resulting model was then applied to the left-out participants. The process was repeated 10 times until all folds served as the test group. Model performance was evaluated as the Pearson correlation between predicted and observed target scores. This pipeline was repeated 1000 iterations to allow different partitioning of folds with the constraint that family members could only be in the same fold. The mean and standard deviation of the model performances across iterations are presented (Fig. 1; see Fig. S4 for the distributions of model performance).

**Figure 1:**
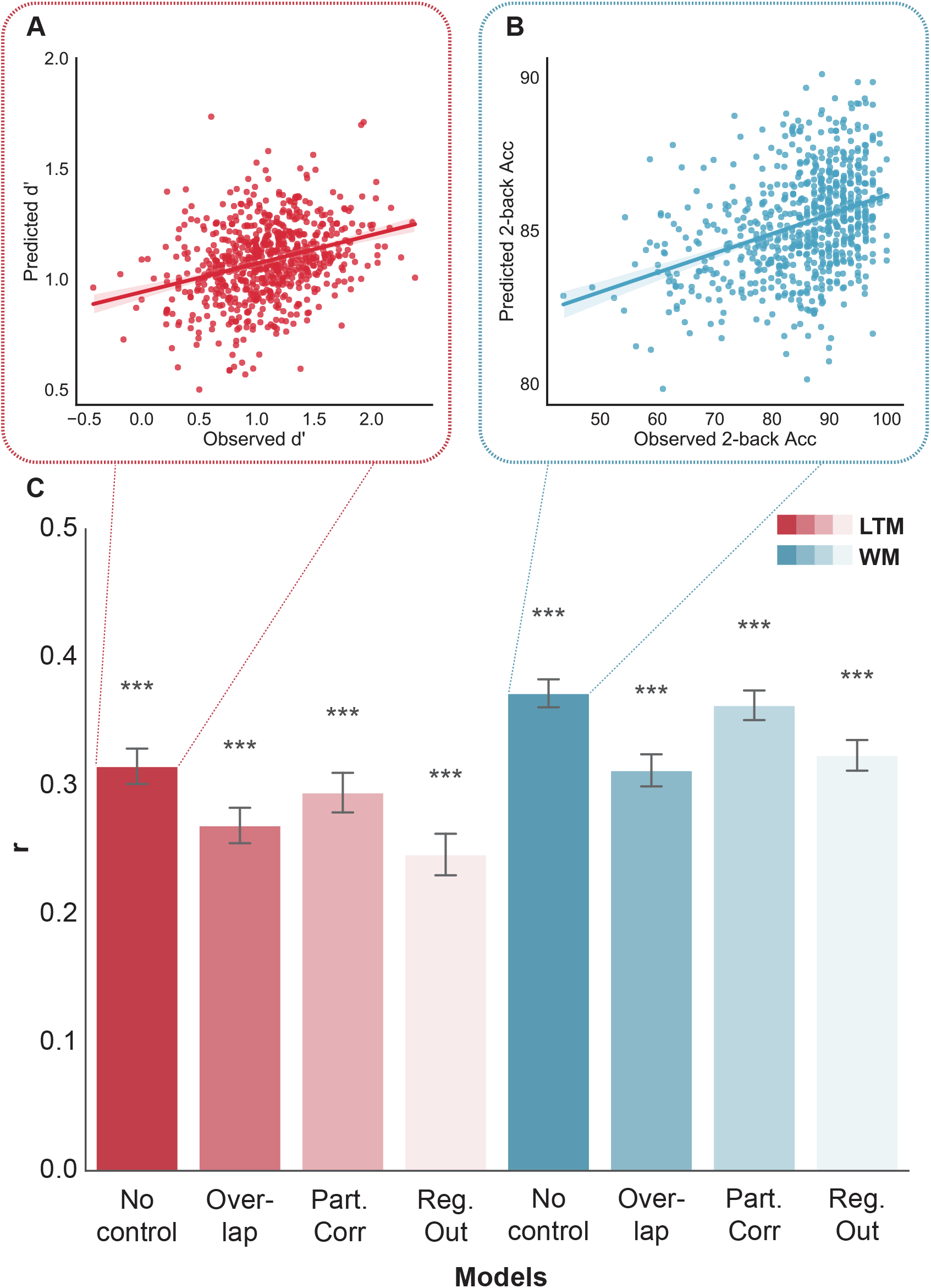
Functional connectivity during n-back tasks robustly predicts long-term memory (LTM) and working memory (WM). (A) A scatterplot depicting the correlation between predicted and observed LTM for one iteration where the *r* value matched the mean model performance across all iterations. (B) A scatterplot depicting the correlation between predicted and observed WM for one iteration where the *r* value matched the mean model performance across all iterations. (C) Summary of model performance for predicting LTM and WM with different model training procedures. No control: full model without controlling for the non-target behavior. Overlap: limiting edge selection to those correlated with both LTM and WM. PartCorr: controlling for the non-target behavior during edge selection using partial correlation. RegOut: building models to predict the residuals in target behavior after regressing out the non-target behavior. Bar height: mean Pearson’s *r* between predicted and observed scores across iterations. Error bars: standard deviation of the *r* values. ***: *p* < .001.

To assess the statistical significance of model performance, we repeated the pipeline above with permuted participant-score correspondence on each iteration (with the same family structure constraint). *P* values were calculated as the fraction of iterations where the unpermuted models performed worse than the best performing null model. All *p* values were corrected for multiple comparisons using the Bonferroni correction.

### 2.1 LTM model has some shared components with, but is also distinct from WM model

We first demonstrated that FC during n-back task successfully predicted out-of-scanner recognition memory performance (Fig. 1A and 1C; mean *r* = 0.32, *p* < .001). Replicating Avery et al.(2019), a separate model built from the same n-back task FC also successfully predicted 2-back accuracy (Fig. 1B and 1C; mean *r* = 0.37, *p* < .001).

As LTM and WM measures were moderately correlated in our sample (*r* = 0.25, 95% CI from 1000 bootstrapping iterations: [0.19. 0.31]), next we investigated the shared and distinct components of brain networks supporting these two functions. To probe the overlap of the two networks, we only included edges that are correlated with both WM and LTM (at *p* < 0.01) during feature selection. The average number of edges included in the model was thus reduced to 373.35, an 88.77% reduction from the 3323.72 edges for the full WM model and an 83.50% reduction from the 2263.35 edges for the full LTM model. Model performances remained robust for both memory constructs even when limiting features to only overlapping edges (Fig. 1C; LTM: mean *r* = 0.27, *p* < .001; WM: mean *r* = 0.31, *p* < .001), suggesting that WM and LTM are supported by partially overlapping brain networks.

After demonstrating that there were shared components between LTM and WM models, we sought to investigate the distinct functional architectures underlying LTM and WM. First, we controlled for the non-target measure by running a partial correlation during the feature selection step and included only edges that were strongly correlated with the target behavior even after controlling for the non-target behavior. Both models remained robust after the control (Fig. 1C; LTM: mean *r* = 0.29, *p* < .001; WM: mean *r* = 0.36, *p* < .001). To further investigate whether partial correlation effectively isolated the two memory measures, we tested if the control models generalized to predict the non-target behavior. Interestingly, both control models still generalized to predict the non-target behavior, albeit to a lesser degree (trained on LTM, tested on WM: mean *r* = 0.21, *p* < .001; trained on WM, tested on LTM: mean *r* = 0.19, *p* < .001). These generalization results suggest that the partial correlation procedure did not necessarily remove all edges that were correlated with the non-target measure. That is, partial correlation only guaranteed that the selected edges were correlated with the residuals in the target behavior after the non-target behavior was regressed out from both the edge strengths and target behaviors. However, this did not guarantee that the selected edges were not correlated with the non-target behavior^2^. Furthermore, the partial correlation models could still be capturing some of the shared components between WM and LTM behavior measures and thus generalize to predict the shared components in the full non-target scores.

To further establish that predictive models are able to capture the unique aspect of each memory measure, we regressed out the shared component between the two measures, built models and tested them on the residuals (in a fully cross-validated manner, see Methods for details). Both models remained robust (Fig. 1C; LTM: mean *r* = 0.25, *p* < .001; WM: mean *r* = 0.33, *p* < .001). Interestingly, while the model trained on only LTM residuals still marginally predicted the full LTM scores (mean *r* = 0.21, *p* = .048, uncorrected), the WM residuals model did not significantly predict the full WM scores (mean *r* = 0.16, *p* = 0.401), demonstrating that the shared variance between WM and LTM explained more of the variance in WM than in LTM. In contrast to the partial correlation models, these models did not generalize to predict the residuals in the non-target behavior (trained on LTM residuals, tested on WM residuals: mean *r* = 0.002, *p* = .985; trained on WM residuals, tested on LTM residuals: mean *r* = 0.13, *p* = .253). Collectively, these results suggest that there were separable networks supporting LTM and WM respectively.

### 2.2 LTM model engages multiple networks and is spatially distinct from WM model

To identify the anatomy of the predictive models, for each predictive model, we first identified the reliable edges that were selected on 100% of the cross-validation iterations (i.e., [1000*×*10=] 10000 iterations in total). Then for each selected edge, we averaged the assigned weights (normalized by the sum of absolute values of ridge regression weights for all edges during a given iteration) across all 10000 iterations. The value assigned to each edge this way can be thought of a predictive utility where the absolute value indicates the fractional contribution of the feature to model performance and the sign represents the relationship between the feature and the target behavior. We first visualized the distribution of these reliable edges by grouping them into macroscale brain regions. We then explored the distributions of the models by grouping the nodes into 10 previously defined canonical networks (Greene et al., 2018; Noble et al., 2017) and averaging the weights of features within each network and between each pair of networks.

We first compared the similarities between the different predictive models for the 2 target behaviors (LTM/WM) *×* 4 procedures (no control, partial correlation, regressing out control behavior, and overlapping edges) by calculating the Spearman rank correlation between each pair of the predictive models on the network level (Fig. 2A). The features identified for the same target behavior are highly similar to each other except when features were forced to be the overlapping edges between the two memory measures. Because we only constrained the set of edges to be included in the ridge regression, the relative weights of the edges could still differ between the two target behaviors if they required distinct configuration of these set of edges. However, instead we observed high rank similarity on the network level (*r* = 1.00) between the overlapping models for WM and LTM, suggesting that the overlapping edges could be supporting cognitive processes that contribute to both the retention of WM and LTM.

**Figure 2:**
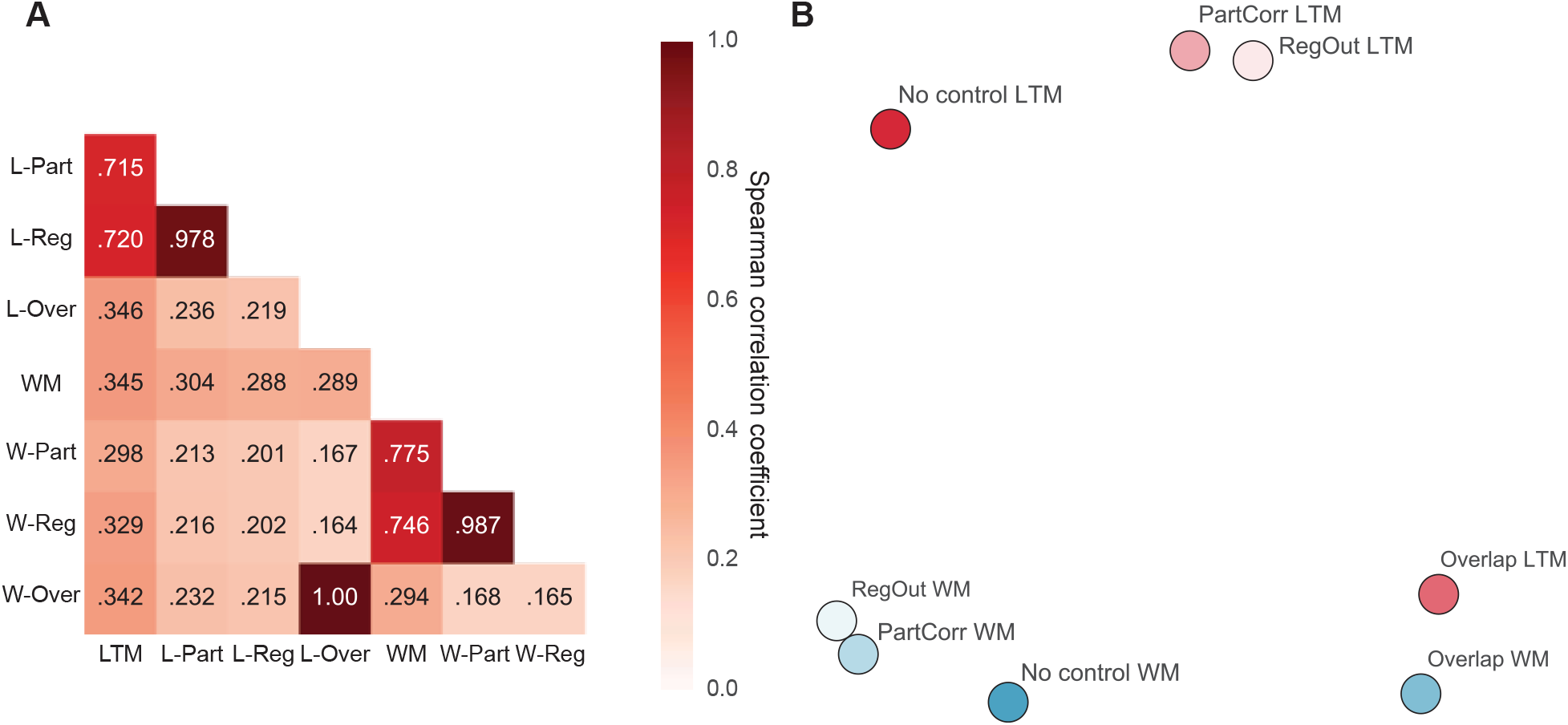
Comparison of the anatomical distribution of the different predictive models. (A) Spearman correlations between the features of different models at the network level. LTM: long-term memory full model; L-Part: LTM partial correlation model; L-Reg: LTM regressing out model; L-Over: LTM overlap model; WM: working memory full model; W-Part: WM partial correlation model; W-Reg: WM regressing out model; W-Over: WM overlap model. (B) Multidimensional scaling plot illustrating the distance between the different models at the network level. Distance was calculated as the Euclidean distance between each pair of models after mean-centering and scaling along each dimension. No control: full model without controlling for the non-target behavior. Overlap: limiting edge selection to those correlated with both LTM and WM. PartCorr: controlling for the non-target behavior during edge selection using partial correlation. RegOut: building models to predict the residuals in target behavior after regressing out the non-target behavior.

As shown in Fig. 2B, the distributions of predictive networks for the same target behavior from the no control, partial correlation and orthogonalizing behavior procedures are quite similar to each other on the canonical network level (LTM: minimum *r* = 0.72; WM: minimum *r* = 0.75). For concision and clarity, only the predictive networks from the regressing out control behavior procedure are reported below (Fig. 3) to demonstrate the patterns because they exhibited the largest distinction between WM and LTM (*r* = 0.20) with the corresponding results from predictive networks from other procedures in Fig. S5. Additionally, we visualize the anatomical distribution in terms of macroscale regions of the overlapping models for both LTM and WM.

**Figure 3:**
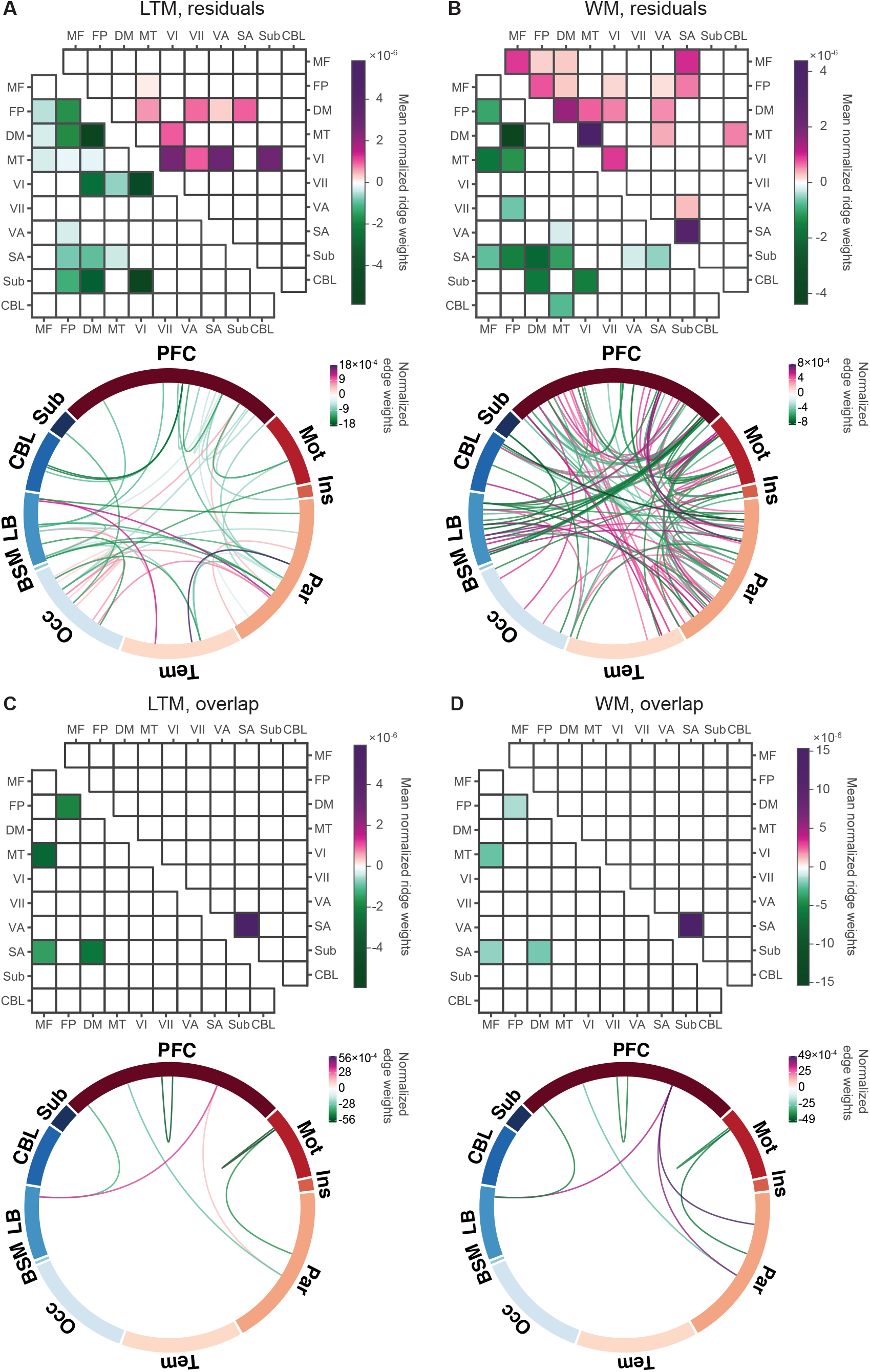
Anatomical distributions of predictive features. For each panel, on the top are results from grouping nodes at the network level. Purple: features with positive weights, green: features with negative weights, shade: relative model contribution. Network labels: MF - medial frontal, FP - frontoparietal, DM - default mode, MT - motor, VI - visual I, VII - visual II, VA - visual association, SA - salience, Sub - subcortical, CBL - cerebellum. On the bottom are results from grouping nodes into macroscale brain regions. The entire circle represents all 368 nodes, divided into the macroscale brain regions. The different locations on each ribbon represent the different nodes that belong to this region. Each line represents a reliable predictive edge. Purple: features with positive weights, green: features with negative weights, shade: relative edge weight. Macroscale regions labels: PFC - prefrontal cortex, Mot - motor cortex, Ins - insula, Par - parietal cortex, Tem - temporal cortex, Occ - occipital cortex, BSM - brainstem, LB – limbic regions (including the cingulate cortex, amygdala and hippocampus), Sub – subcortical regions (including the thalamus and striatum), CBL - cerebellum. (A) Predictive features of the model trained for the residuals in LTM after regressing out WM. (B) Predictive features of the model trained for the residuals in WM after regressing out LTM. (C) Predictive features of the model trained for LTM (without regressing out WM) with only edges correlated with both WM and LTM. (D) Predictive features of the model trained for WM (without regressing out LTM) with only edges correlated with both WM and LTM.

For the LTM model (Fig. 3A), edges in the Visual I, Visual I-Visual Association and Visual I-Subcortical networks emerged as the most important positive features whereas edges in the Default Mode, Visual I and Visual I-Subcortical networks as the most important negative features. The importance of edges in the visual networks was further demonstrated by the distribution of predictive edges across the macroscale brain regions as the occipital lobe was involved in most of the predictive features. In contrast to the LTM model, the WM model contained edges that were more widespread (Fig. 3A bottom vs. Fig. 3B bottom) and engaged more networks (Fig. 3B). Edges in the Motor, Visual I and Salience networks emerged as the most important positive features in predicting WM whereas edges in the Default Mode-Frontoparietal, Default Mode-Salience and Visual I-Subcortical networks emerged as the most important negative features. When we limited the features to only edges correlated with both WM and LTM (Fig. 3C and 3D), the predictive models for the two measures were highly similar both in terms of the distribution across macroscale brain regions and canonical networks. Edges within the prefrontal and parietal regions of the Salience network were the most important positive features whereas edges between the Medial Frontal and Motor networks were the most important negative features.

The different patterns of critical features between LTM and WM networks are further supported by the following two complementary analyses. First, we only preserved connections within a given network during the feature selection step. For the full LTM scores, preserving the Visual I, Default Mode, and Salience networks led to best prediction performance (Fig. 4A) whereas for the full WM scores, preserving the Frontoparietal, Salience, or Motor networks lead to the best prediction performance (Fig. 4B). Secondly, we virtually ‘lesioned’ each of the 10 networks by excluding all connections within and from the network during the feature selection step. Mirroring the results above, the lesioning analyses again highlighted the importance of Visual I and Default Mode networks to LTM (Fig. 4C) and that of the Motor, Frontoparietal and Salience networks to WM (Fig. 4D).

**Figure 4:**
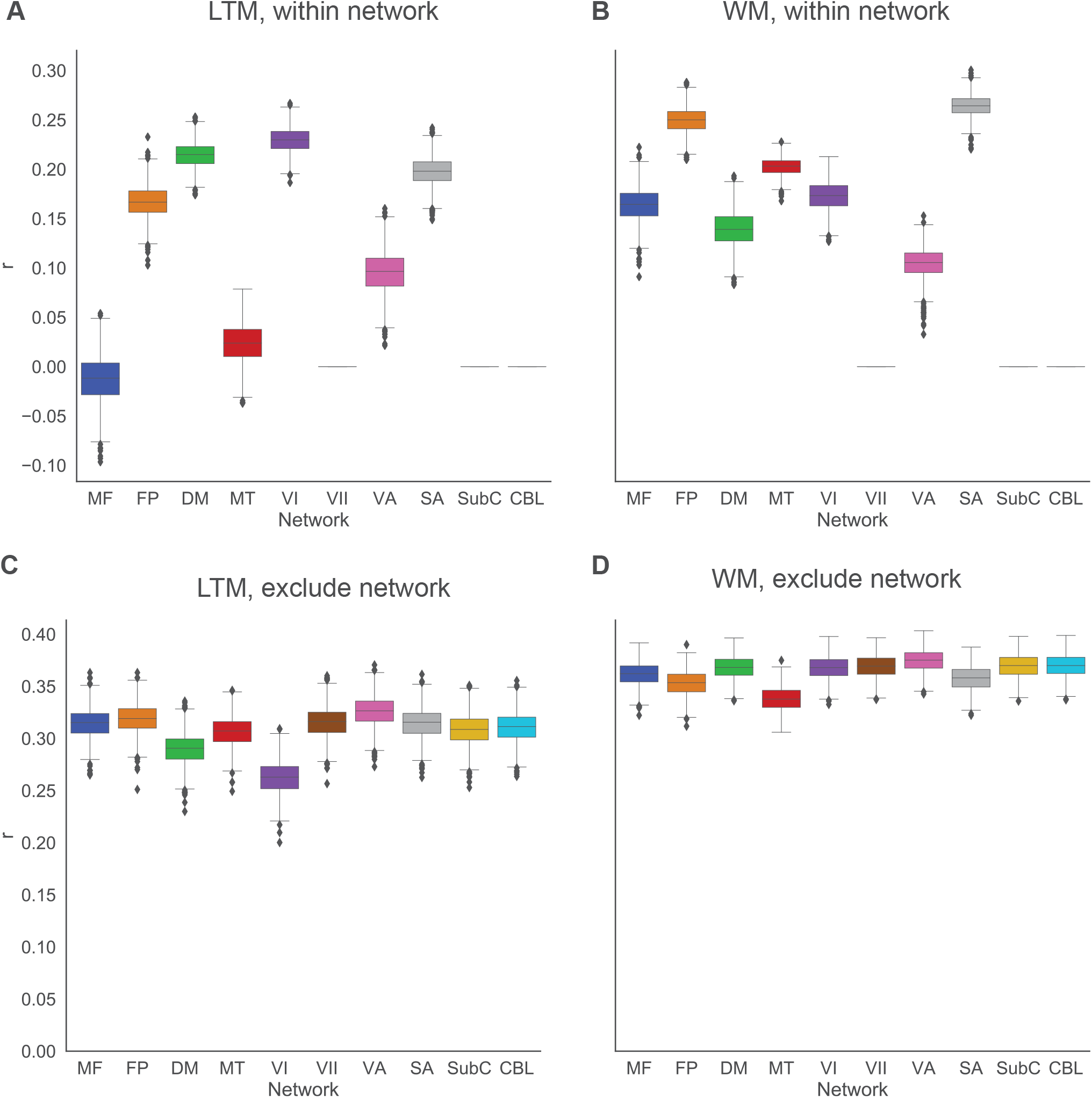
Boxplots of model performance when controlling for each of the ten different networks. Network labels: MF - medial frontal, FP - frontoparietal, DM - default mode, MT - motor, VI - visual I, VII - visual II, VA - visual association, SA - salience, SubC - subcortical, CBL - cerebellum. For some networks, no edges passed the thresholding step (*p* < 0.01) and thus no prediction was generated. This is reflected by missing boxes. (A) Predicting LTM when edge selection was restricted to within-network edges. (B) Predicting WM when edge selection was restricted to within-network edges. (C) Predicting LTM when edge selection was performed after excluding edges from a given network. (D) Predicting WM when edge selection was performed after excluding edges from a given network.

### 2.3 ROIs previously associated with LTM are useful but not sufficient to predict LTM

Although the anatomical distribution analyses above suggest that the predictive network of individual differences in LTM engages multiple functional networks across the whole brain, in the following section, we explicitly related our findings to the brain regions identified in the previous literature to be important for LTM formation. To this end, we first defined an anatomical medial temporal lobe mask based on the Harvard-Oxford Atlas and then defined a functional mask related to subsequent memory effect using the NeuroSynth database (Yarkoni et al., 2011).

We identified 11 nodes in the Shen atlas that overlap with the anatomical medial temporal lobe mask over 5% (see Fig.5A for a visualization of these nodes) and investigated the role of the medial temporal lobe during model training (see Cognitive Prediction in Methods) in four ways: 1) including only edges within these 11 nodes; 2) including only edges connecting within these nodes and from these nodes to other nodes; 3) including only edges connecting the medial temporal lobe nodes and nodes outside of the medial temporal lobe (but not within the medial temporal lobe); 4) excluding any edges that involve medial temporal lobe nodes. The statistical significance of each model was assessed via permutation tests. The performances of different unpermuted models were compared using Wilcoxon signed-rank tests. All *p* values were corrected for multiple comparisons using Bonferroni correction.

The results from medial temporal lobe analysis were shown in Fig. 5C. Both LTM and WM models built from edges within medial temporal lobe did not provide significant prediction of the target behavior. All other models performed above chance (all *p*s ≤ .05). These results exhibited two salient patterns: firstly, medial temporal lobe is more useful for predicting individual differences in LTM than WM. Limiting feature selection to only medial temporal lobe and medial temporal lobe related edges or only medial temporal lobe related edges led to better prediction of LTM than WM (both *p*s < .001), consistent with the previous finding that medial temporal lobe is more important for retaining information over longer timescale (Eichenbaum et al., 2007). Secondly, medial temporal lobe alone is not sufficient for predicting LTM. LTM models with just medial temporal lobe or medial temporal lobe related edges performed worse than the full LTM model (only medial temporal lobe and medial temporal lobe related edges: mean Δ*r* = 0.09, *z* = 38.72, *p* < .001; only medial temporal lobe related edges: mean Δr = 0.09, *z* = 38.72, *p* < .001). Furthermore, the performance of the LTM model built from only edges connecting the medial temporal lobe and other regions was actually a little bit better than that of the LTM model built from both edges within the medial temporal lobe and those connecting the medial temporal lobe and other regions (mean Δ*r* = 0.001, *z* = 2.32, *p* = .020), suggesting that edges within medial temporal lobe did not provide much additional predictive power. Lastly, excluding any edges involving medial temporal lobe led to only a statistically significant but small hit in prediction performance compared to the full LTM model (mean Δr = 0.01, *z* = 19.54, *p* < .001), further suggesting that individual differences in LTM encoding are related to a whole-brain network, rather than just localized areas.

**Figure 5:**
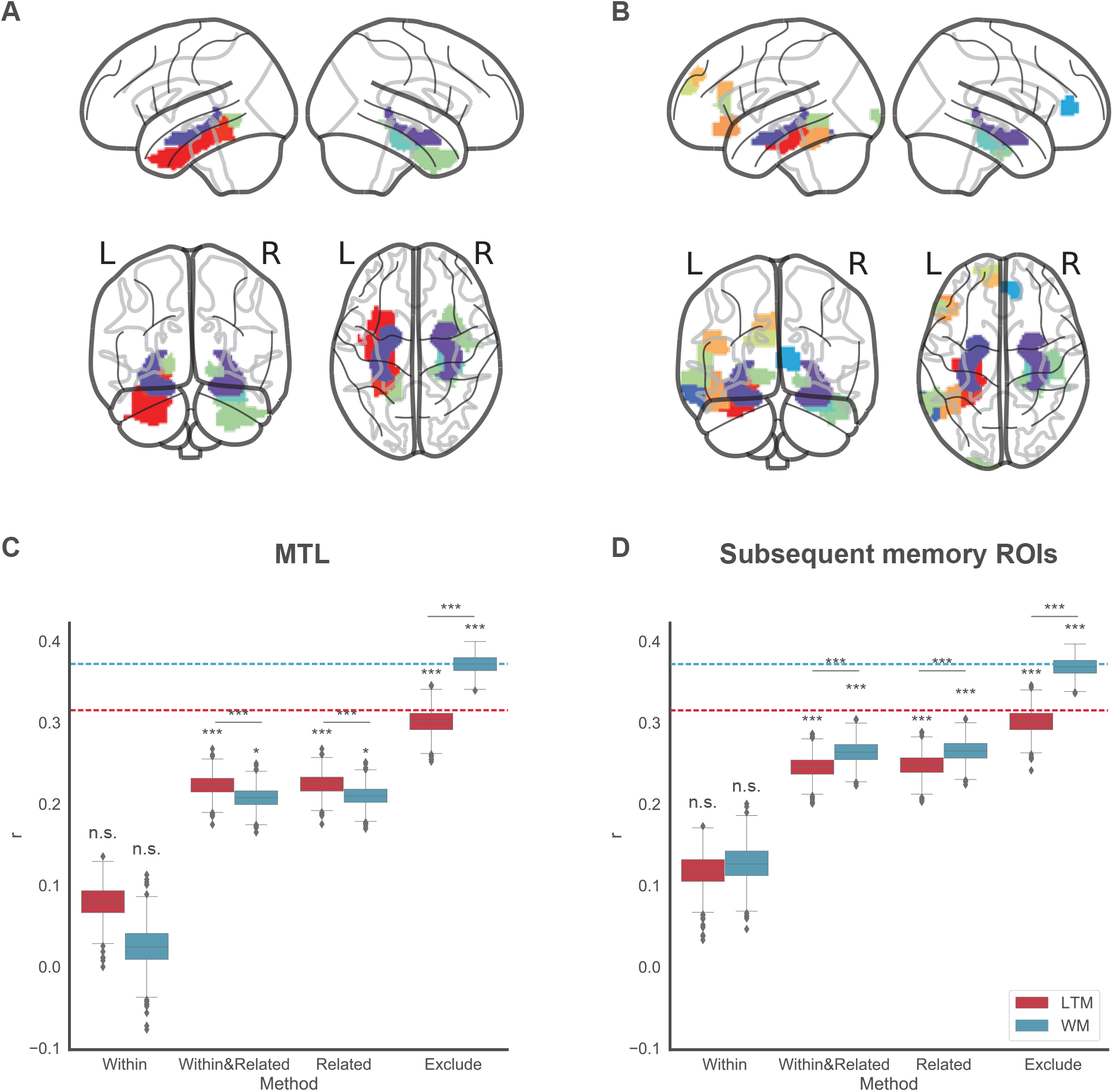
Results from controlling for ROIs previously associated with LTM. (A) Nodes in the Shen atlas that overlap with the medial temporal lobe (MTL) mask over 5%. (B) Nodes in the Shen atlas that overlap with the subsequent memory mask over 5%. Individual nodes are represented by distinct colors. (C) Boxplot of model performance when controlling for nodes in MTL using different methods. (B) Boxplot of model performance when controlling for ROIs previously associated with subsequent memory effect using different methods. Within: limiting edges to only those within the control ROI, WithinRelated: limiting edges to those within and from the control ROI, Related: limiting edges to only those between nodes in the control ROI and nodes outside of the control ROI, Exclude: excluding any edges involving nodes in the control ROI. Red: LTM, blue: WM. Dash lines represent the mean performance of the LTM/WM models without any control. *: *p* < .05, **: *p* < .01, ***: *p* < .001.

Next, we explored the role of regions showing subsequent memory effect in previous literature with the same approaches used for the medial temporal lobe analysis. There were 19 nodes in the Shen atlas that overlap with the subsequent memory mask more than 5% (see Fig.5B for a visualization of these nodes). Note that the subsequent memory mask includes regions beyond the medial temporal lobe, in the frontal and occipital lobes.

The results from subsequent memory ROIs analysis were shown in Fig. 5D. Both LTM and WM models built from edges within the subsequent memory mask did not provide significant prediction of the target behavior. All other models performed above chance (all *p*s < .001). In contrast to what we observed above for the medial temporal lobe mask, for the subsequent memory ROIs, we did not observe an advantage of LTM. On the contrary, when feature selection was limited to only edges within and connecting the subsequent memory mask or only edges between the mask and other regions, the WM models performed better than LTM models (both *p*s < .001), potentially because of the inclusion of additional brain regions in the frontal lobe. Similar to what we observed in the medial temporal lobe analyses, although the subsequent memory network is useful for predicting LTM, it’s not sufficient. Both the LTM model with just edges within and connecting the subsequent memory ROIs and one with just edge connecting the subsequent memory ROIs and other nodes performed worse than the full LTM model (within and related edges: mean Δr = 0.07, z = 38.69, *p* < .001; only related edges: mean Δr = 0.07, *z* = 38.68, *p* < .001). This is further supported by the observation that including edges both within and from subsequent memory ROIs led to slightly worse performance in predicting LTM scores than including only those connecting these ROIs and other regions (mean Δr = 0.003, *z* = 4.26, *p* < .001) and that excluding any edges involving the subsequent memory ROIs still yielded a robust prediction of LTM compared to the full model (mean Δr = 0.01, *z* = 19.56, *p* < .001).

## 3 Discussion

We built a predictive model of individual differences in LTM performance based on whole-brain FC during an n-back task. By comparing the LTM model and a separate model built to predict WM performance based on the same FC data, we found that there are some shared components between WM and LTM but more critically, these two networks are also dissociable from each other, shedding new light on the distinction between the neural networks supporting WM and LTM. In addition, we also showed that models built on ROIs previously shown to be associated with within-subject fluctuations in LTM performance are useful but not sufficient to capture across-subject differences in LTM, demonstrating the importance of considering large-scale networks in understanding the functional architecture supporting LTM.

Our approach complements previous research on the relationship between LTM and WM in two important ways. First, we were able to build LTM and WM models with only features that were predictive of both WM and LTM, suggesting that they are supported by overlapping edges mainly in the salience network. Furthermore, the relative importance of these selected edges was highly similar for predicting WM and LTM, further demonstrating that there are shared cognitive processes supporting the retention of information in both short- and long-timescales. Second, we showed that the LTM model remained robust when we controlled for WM and vice versa. Given that the LTM and WM models were identified from the same brain activities during the n-back tasks and thus effects from stimuli and neural states were held the same for the two models, our finding of dissociable WM and LTM networks revealed a true distinction between the neural architectures supporting the retention of the same information in WM vs. LTM. Functional connections in the visual, default mode and subcortical networks contributed more to the prediction of LTM, whereas those in the motor, frontoparietal, default mode and salience networks were more important for the prediction of WM.

The partitioning of shared and unique components between LTM and WM networks in our study could be especially useful in identifying which of these components are selectively associated with normal and pathological aging, thus providing implications for developing more targeted treatment. Both WM and LTM showed substantial decline with advanced aging (Park et al., 1996; Park et al., 2002). However, memory systems are differentially affected by normal aging and age-related disease such as Alzheimer’s Disease (Gabrieli, 1996). Moreover, previous work has suggested that there are multiple factors linked to memory decline in aging and distinguished between changes in the frontal-striatal system, which primarily affects executive functioning, and changes in the medial temporal lobe memory system (Buckner, 2004). Separate studies have found that functional connectivity measured during resting state is predictive of WM (Avery et al., 2019), LTM (Meskaldji et al., 2016) and general cognitive impairment (Lin et al., 2018) in an aging population. These studies suggest that there were functional connections shared by models of these memory-related constructs, such as those in the frontoparietal control and default mode networks. However, there were also differences: the motor network has been implicated in the WM and general cognitive impairment networks, whereas the thalamus plays an important role in modulating LTM. Extending on these previous results, our study investigated the overlapping and unique features supporting LTM and WM in the same sample and thus better controlled for the effect of samples and analysis pipelines on the identified features. The overlapping and unique features we found can complement behavioral testing and serve as a tool for characterizing distinct neural profiles of cognitive impairment due to normal aging or pathological conditions.

In addition to providing insights into the functional architectures supporting WM and LTM, our study also poses interesting questions about the neural sources of between- and within-individual differences in LTM performance. We found that although ROIs previously found to be critical to within-individual fluctuations in LTM encoding in group-level analyses captured some variance in LTM performance across individuals, the predictive power of a model built on only these ROIs was substantially weaker than when we considered all the possible edges across the whole brain. Furthermore, excluding these ROIs from our analyses led to only a negligible drop in predictive power. There are at least two possible explanations for this pattern of results. First, these LTM ROIs were originally derived from group-based studies where data from individual subjects were aggregated to find common neural processes underlying LTM (for reviews, see Paller and Wagner, 2002; Squire et al., 2004). Therefore, these ROIs should show stability across individuals and may not be informative for revealing between-subject variance. Second, these LTM ROIs were identified using task-activation whereas the neural measure in our study is functional connectivity. These two measures are not redundant: in fact, a certain brain region can show decrease in activity but increase in connectivity with other brain regions and vice versa (Bassett et al., 2015). Consistent with recent works in other cognitive domains (Bassett & Sporns, 2017; Bullmore & Sporns, 2009; Petersen & Sporns, 2015; Shine et al., 2019; Turk-Browne, 2013), our results show that the ability to successfully encode information into LTM is an emergent property of large-scale functional brain networks, rather than localized brain regions.

Our finding that the LTM model extends beyond ROIs identified in previous studies is also consistent with the recent findings by Keerativittayayut et al.(2018), who studied how dynamic FC during encoding captured within-individual fluctuations in memory performance using a classic incidental encoding task and a surprise recognition memory test. They found that within the subsequent memory ROIs, only FC between hippocampus and occipital cortex differed between good and bad encoding states. Furthermore, they also observed that edges in areas beyond the subsequent memory ROIs, mostly in the salience, subcortical, default mode and visual networks, played important role in distinguishing encoding states. Corroborating these within-individual findings, our results highlighted the importance of these same networks in predicting between-individual differences. Recent work showed that a sustained attention model built to predict attention performance across individuals generalized to predict the temporal fluctuations of attention within the same individual, demonstrating that the FC-based sustained attention model captures a combination of trait-like and state-like sustained attention (Rosenberg et al., 2020). The similarities in the LTM-related features identified by Keerativittayayut et al.(2018) and our study suggest that the LTM network in our study may also contain both features that underlie trait-level differences in LTM and those that underlie state-level fluctuations. Future work can directly compare the within- and between-subject LTM networks to delineate the neural sources of within- and between-individual differences in LTM performance.

The LTM predictive network in the current study was identified during encoding. However, successful memory performance also depends on post-encoding processes such as consolidation and retrieval. Previous work found that FC between the hippocampus and sensory cortex during the rest period after encoding were correlated with later associative memory performance above and beyond FC during encoding (Tambini & Davachi, 2019; Tambini et al., 2010). A similar set of brain regions that are related to better encoding are also implicated in successful retrieval (Rugg & Vilberg, 2013). Our individual differences approach can be applied to identify consolidation and retrieval networks to reveal how these functional networks differ or are related to each other.

In sum, our study demonstrates the utility of adopting an individual differences approach in revealing novel aspects of the neural architecture supporting the formation of LTM and how it is related to and separable from other memory measures or processes, using comparison with WM as a case study. Identifying predictive models of different cognitive components involved in LTM not only provides new insights into the interaction and distinction of these components but also illuminates the development of more specific biomarkers targeting the separable components in memory-related disorders.

## Acknowledgements

Data were provided by the Human Connectome Project, WU-Minn Consortium (Principal Investigators: David Van Essen and Kamil Ugurbil; 1U54MH091657) funded by the 16 NIH Institutes and Centers that support the NIH Blueprint for Neuroscience Research, and by the McDonnell Center for Systems Neuroscience at Washington University. This work was supported by NIH Grant MH108591 to M.M.C.

## Methods

### Dataset

Data used in this study were released as part of the Human Connectome Project (HCP) S1200 release, described below.

#### Participants

We restricted our analyses to those participants who completed the two n-back fMRI runs and both runs of the recognition memory conducted outside of the scanner. Additionally, we excluded participants whose motion during the n-back runs did not meet any of following criteria: 1) grand mean root mean square (RMS) relative motion across both runs < 0.1 mm; 2) maximum mean RMS relative motion < 0.16; 3) maximum translation < 2 mm; 4) maximum rotation < 3^*°*^. These motion thresholds were selected based on previous related work to mitigate the effect of excessive head motion on FC (e.g., Greene et al., 2020; Rosenberg et al., 2015). 708 participants were included in the final analysis (371 females, ages 22 to 36 years [*M* = 28.51, *SD* = 3.70, Median = 29]).

#### N-back working memory tasks

Details of the task can be found in Barch et al.(2013). Briefly, participants performed two runs of the n-back tasks while being scanned (about 5 minutes each). Each run consisted of eight task blocks of 10 trials. Within each run, half of the blocks are 0-back and the other half are 2-back. During each block, images can be from one of the four categories: faces, places, tools and body parts. 2-back accuracies (% correct; collapsed across all four stimulus categories) were chosen as the target WM measure (*M* = 85.23, *SD* = 9.74, range = [43.58, 100]).

#### Recognition memory task

After the participants completed all fMRI runs during the session with the WM tasks, they were given a “Remember, Know, New” item recognition test for the faces and places images (48 in total) presented during the WM tasks, with an equal number of new faces and places. For each participant, d’ was calculated as z(hit rate) – z(false alarm rate) using the stats.norm.ppf function from the scipy package in Python. Note that we did not distinguish between ‘Remember’ and ‘Know’ responses and counted both as ‘Old’ responses. d’ was used to measure long-term memory performance (*M* = 1.08, *SD* = 0.41, range = [-0.42, 2.38]).

#### Imaging parameters and preprocessing

Details of imaging parameters (Essen et al., 2013) and preprocessing pipelines (Glasser et al., 2013) have been published elsewhere. Briefly, all fMRI data were acquired on a 3T Siemens Skyra using a slice-accelerated, multiband, gradient-echo, echo planar imaging (EPI) sequence (TR = 720 ms, TE = 33.1 ms, flip angle = 52 degrees, resolution = 2.0 mm3, multiband factor = 8). Data from the two WM runs (one left/right [LR] phase encoding run and one right/left [RL] phase encoding run) were used for this work. The scanning protocol (as well as procedures for obtaining informed consent from all participants) was approved by the Institutional Review Board at Washington University in St. Louis. Use of HCP data for these analyses was deemed exempt from IRB review by the Yale Human Investigation Committee. The HCP minimal preprocessing pipeline was used on these data, which includes artifact removal, motion correction, and registration to standard space(Glasser et al., 2013). All subsequent preprocessing steps were similar to the preprocessing procedures in previous CPM studies (e.g., Finn et al., 2015; Rosenberg et al., 2015) and were conducted with FSL(Jenkinson et al., 2011), FreeSurfer (Fischl, 2012)and the Python module Nilearn(Abraham et al., 2014), which includes detrending, regressing out covariates of no interest (mean signals from gray matter, white matter and cerebrospinal fluid and 24 motion regressors [6 motion parameters, 6 temporal derivatives, and their squares]) and bandpass filtering (range: [0.01 – 0.1 Hz]). Mean frame-to-frame displacement (FD) was calculated following (Power et al., 2014) for each run and averaged to yield a motion value per each included subject to be used for motion control analysis (e.g., partial correlation-based feature selection; see below). All subsequent analyses and visualization were performed with Matlab (Mathworks) and Python.

### Functional parcellation

The Shen 368-node atlas (Shen et al., 2013) was applied to the preprocessed HCP data in standard space. After parcellating the data from each run into 368 nodes, the mean time courses of each node pair were correlated using Pearson’s *r* and correlation coefficients were Fisher’s z transformed, generating two 368*×*368 connectivity matrices per participant. These two matrices were then averaged to produce one matrix per participant.

### Cognitive prediction

A modified version of connectome-based predictive modeling (CPM; Finn et al., 2015; Rosenberg et al., 2015; Shen et al., 2017) was used to predict the target behavior (2-back accuracy or d’) from the connectivity matrices using ridge regression (Gao et al., 2019; Greene et al., 2020). The models were trained and tested in a nested 10-fold cross-validated manner.

First, the participants were split into ten groups, respecting family structure such that family members were always assigned to the same group. Nine of these groups served as training data; in the training set, features were selected on the basis of their Pearson correlation with the target score with a threshold of *p* < 0.01 (we tested a wide range of *p* values to make sure the results were consistent; see Fig. S1). Selected edges were then submitted as predictors (with the target score as response) to an L2-constrained linear least squares regression (lasso function in Matlab with Alpha set to 1e-6). The regression coefficients and intercept for each regression were estimated with an inner 10-fold cross-validation loop using only the training data. The set of parameters with the minimum cross-validated mean squared error (MSE) was selected. These fitted coefficients and intercept were then applied to the corresponding edges in the left-out test participants to predict their target scores. For each fold, model performance was evaluated as the Pearson’s *r* correlation between predicted and observed scores in the test participants. (We also evaluated model performance with *q*^2^ [also called prediction *R*^2^; Scheinost et al., 2019] where 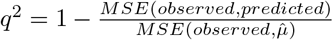 and 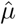 is the mean of the target scores in the training sample [see Fig. S2]. We mainly reported the results from Pearson’s r, and the two measures showed the same pattern.) The above steps were performed iteratively with each group left out once. The resulting *r* values between predicted and observed scores from the 10 folds were averaged to get a measurement of the model performance for the current iteration.

The entire pipeline described above was repeated 1000 times with different group partitions, resulting in 1000 mean *r* values. Significance of the model performance was assessed with 1000 iterations of nonparametric permutation testing (Scheinost et al., 2019) and *p* values were calculated as the fraction of non-permuted iterations on which the mean *r* values were less than or equal to the performance of the best performing null model for the given target score. Resulting *p* values were Bonferroni corrected for multiple comparisons (for the no-control and overlapping models: *α* = 0.05/2 = 0.025; for the partial correlation model: *α* = 0.05/4 = 0.0125; six for the orthogonalization models: *α* = 0.05/6 = 0.008).

### Prediction with only overlapping edges

To investigate the commonalities in the functional networks supporting WM and LTM, we also limited feature selection to only edges that were correlated with both WM and LTM at a threshold of *p* < 0.01 and built ridge regression models to predict the given target measure. Statistical significance was assessed with the same permutation procedure as described above.

### Prediction while controlling for non-target behavior

To investigate whether the functional networks supporting WM and LTM are distinct from each other, we used the following three analyses to control for the non-target behavior: partial correlation, orthogonalization of behavior measures and edge exclusion. Statistical significance was assessed with permuted behavior labels (preserving behavior pairs, i.e., although assigned to the wrong participant, the WM and LTM measures were still from the same participant) but the same analysis pipeline.

#### Partial correlation

We controlled for the non-target measure by running a partial correlation during the feature selection step described in the Cognitive Prediction section. Only edges that were correlated with the target behavior at *p* < 0.01 even after controlling for the non-target behavior were included in the subsequent steps. Additionally, we also tested if these control models trained on one target behavior generalized to predict the non-target behavior (also in a fully cross-validated manner).

#### Orthogonalization of behavior measures

In addition to controlling for the non-target behavior during edge selection, we also conducted a control analysis where we built models to predict the unique aspect of each memory measure by regressing out the shared components between the models. For each fold, before feature selection, using only the training data, we ran a linear regression where the target behavior was the response and the non-target behavior was the predictor. Subsequently, the model was trained to predict the residuals of the target behavior after regressing out the non-target behavior. The intercept and coefficient estimated from the training data were also applied to the two behavior scores in the test set to estimate the residuals in the target behavior after regressing out the non-target behavior. The model performance was thus evaluated as the correlation between the predicted residuals and the actual residuals in the test set. To evaluate how much variance in the full memory scores was captured by the predicted residuals, we also tested the models trained using residuals on the full LTM/WM scores. Additionally, we tested if these control models trained on one target behavior generalized to predict the residuals in the non-target behavior after regressing out the target behavior with the intercept and coefficients estimated from the training set.

#### Edge exclusion (not reported in the main text)

We controlled for the non-target measure by running two Pearson’s correlations with the edges during the feature selection stage: one with the target behavior and the other with the non-target behavior. Thresholding at *p* < 0.01, we obtained two sets of edges that were correlated with the two behaviors. Only edges that were included in the target behavior set but not in the non-target behavior set were selected for the subsequent steps.

Because the model performance from partial correlation and edge exclusion were very similar, to reduce redundancy, we only reported the results from partial correlation and orthogonalization of behavioral measures in the main text. Results of edge exclusion can be found in Fig. S6.

### Evaluating and visualizing feature contributions to a predictive model

To evaluate the contributions of individual features and networks to the prediction of WM and LTM, for each predictive model, we first identified the reliable edges that were selected on 100% of the feature selections (i.e., 10 folds*×*1000 iterations = 10000 feature selections). The contribution of each selected feature was calculated as the mean of normalized beta weight assigned across all iterations of ridge regression. The normalized beta weight for a given edge during an iteration was calculated as the beta weight assigned to this edge divided by the sum of absolute values of ridge regression weights during the given iteration. The value assigned to each edge this way can be thought of a predictive utility where the absolute value indicates the fractional contribution of the feature to model performance, and the sign represents the relationship between the feature and the target behavior.

We first visualized the distribution of these reliable edges by grouping them in to macroscale brain regions. We then explored the distributions of the models by grouping the nodes into 10 previously defined canonical networks (Greene et al., 2020; Noble et al., 2017) and summing the weights of features within each network and between each pair of networks. This sum is then normalized by the total number of pf edges between the given networks to account for differences in network size, resulting in a measure of the mean contribution of an edge in the given network pair.

To further characterize the contributions of different networks, we performed the following two complementary analyses. First, we only preserved connections within a given network during the feature selection step. Secondly, we virtually ‘lesioned’ each of the 10 networks by excluding all connections within and from the network during the feature selection step. Subsequent steps were the same as described in the Cognitive Prediction section.

### The effect of head motion

While we tried to mitigate the effects of head motion on FC by setting stringent inclusion criteria, in this section, we sought to more carefully control for motion in the analysis pipeline. 2-back accuracy was weakly negatively correlated with motion (measured with mean FD averaged across the two WM runs; *r* = -0.09, 95% CI from 1000 bootstrapping iterations: [-0.17. -0.01]) whereas d’ was not correlated with motion (*r* = -0.02, 95% CI: [-0.09, 0.05]). Using the same partial correlation procedure as described above in the *Prediction while controlling for non-target behavior* section, we included mean FD as the covariate during edge selection and repeated the prediction analyses. As an additional way to further control for motion, we also controlled for motion by excluding any edges that correlated with mean FD. Namely, during edge selection, we ran two Pearson’s correlations with the edges during the feature selection stage: one with the target behavior and the other with mean FD. Thresholding at *p* < 0.01, we obtained two sets of edges that were correlated with the memory behavior and motion, respectively. Only edges that were included in the memory behavior set but not in the motion set were selected for the subsequent steps. In both cases, model performance remained robust (see Fig. S3).

---

**Figure S1:**
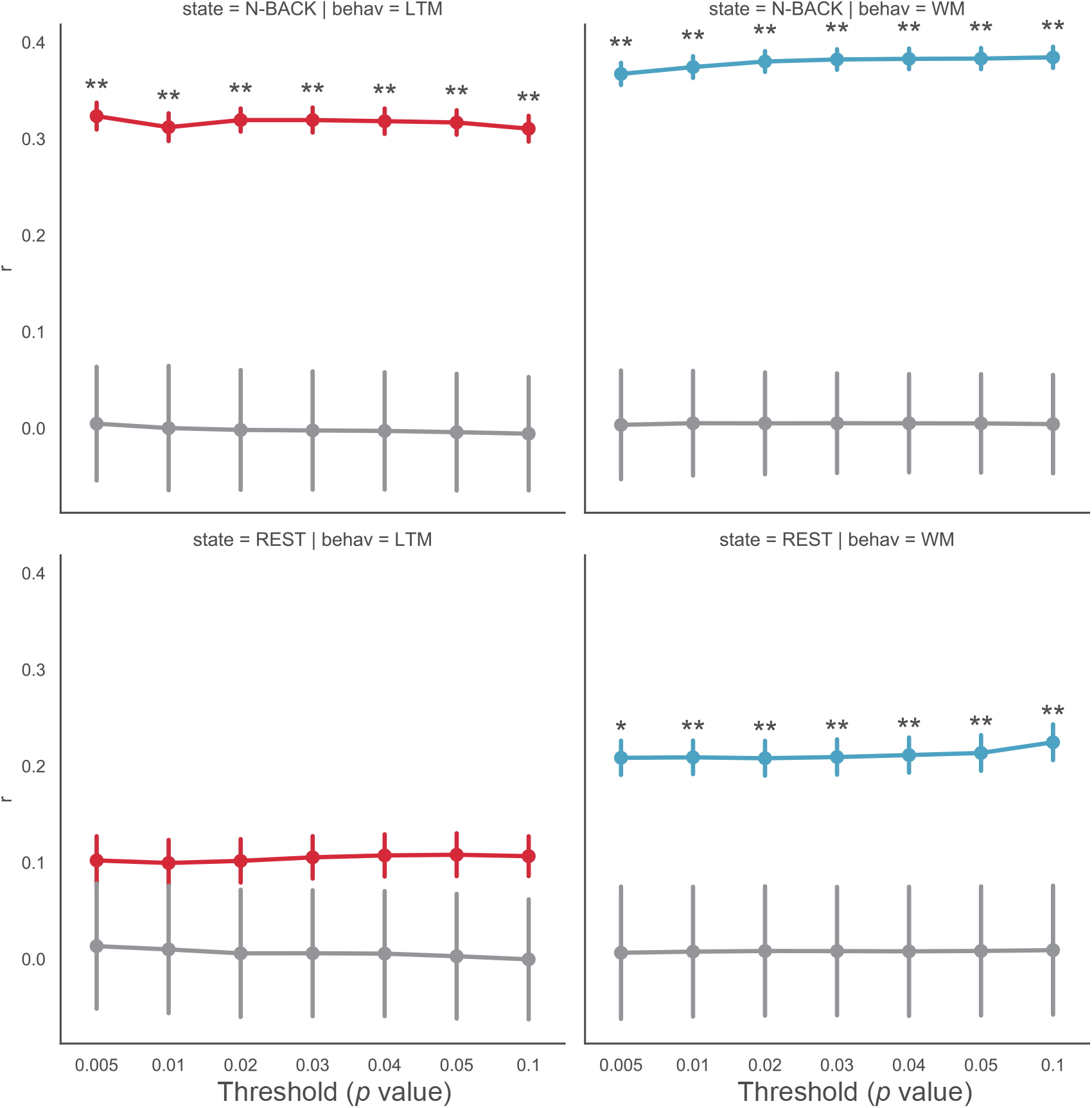
Summary of model performance (measured with Pearson’s r) for predicting long-term memory (LTM; left column) and working memory (WM; right rolumn) using n-back task (top row) and rest (bottom row) fMRI data with different edge selection thresholds. Colored lines: unpermuted model performance, gray lines: permuted model performance. Error bars: standard deviation of the *r* values. The number of iterations was reduced from 1000 to 100 for these analyses to reduce the computational resources used. **: *p* < .01, *: *p* < .05.

**Figure S2:**
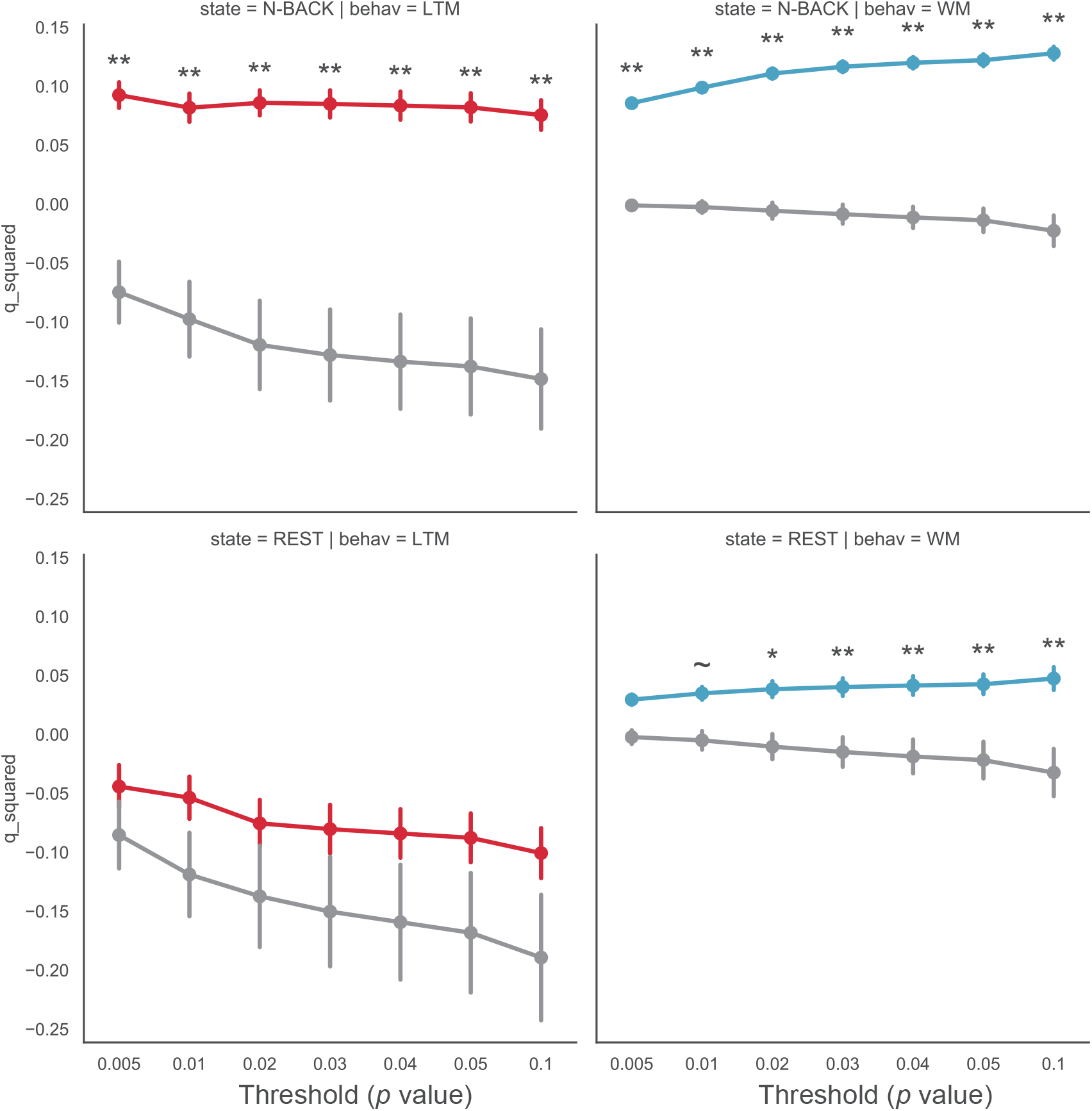
Summary of model performance (measured with q2) for predicting long-term memory (LTM; left column) and working memory (WM; right rolumn) using n-back task (top row) and rest (bottom row) fMRI data with different edge selection thresholds. Colored lines: unpermuted model performance, gray lines: permuted model performance. Error bars: standard deviation of the *r* values. The number of iterations was reduced from 1000 to 100 for these analyses to reduce the computational resources used. **: *p* < .01, *: *p* < .05, *∼*: *p* < .1.

**Figure S3:**
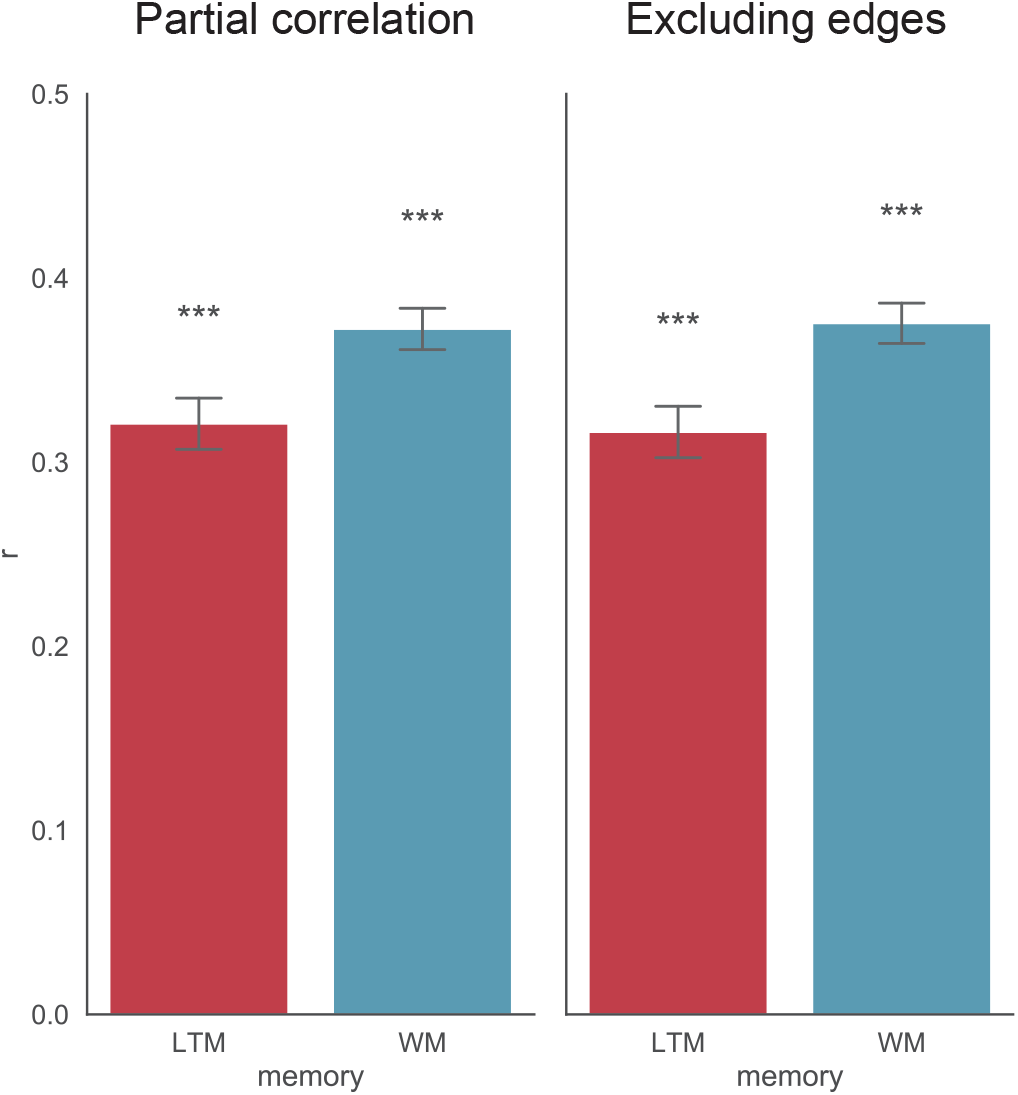
Summary of model performance for predicting LTM and WM after controlling for mean frame-to-frame displacement during edge selection with two different methods (partial correlation and edge exclusion, see methods for more details). ***: *p* < .001

**Figure S4:**
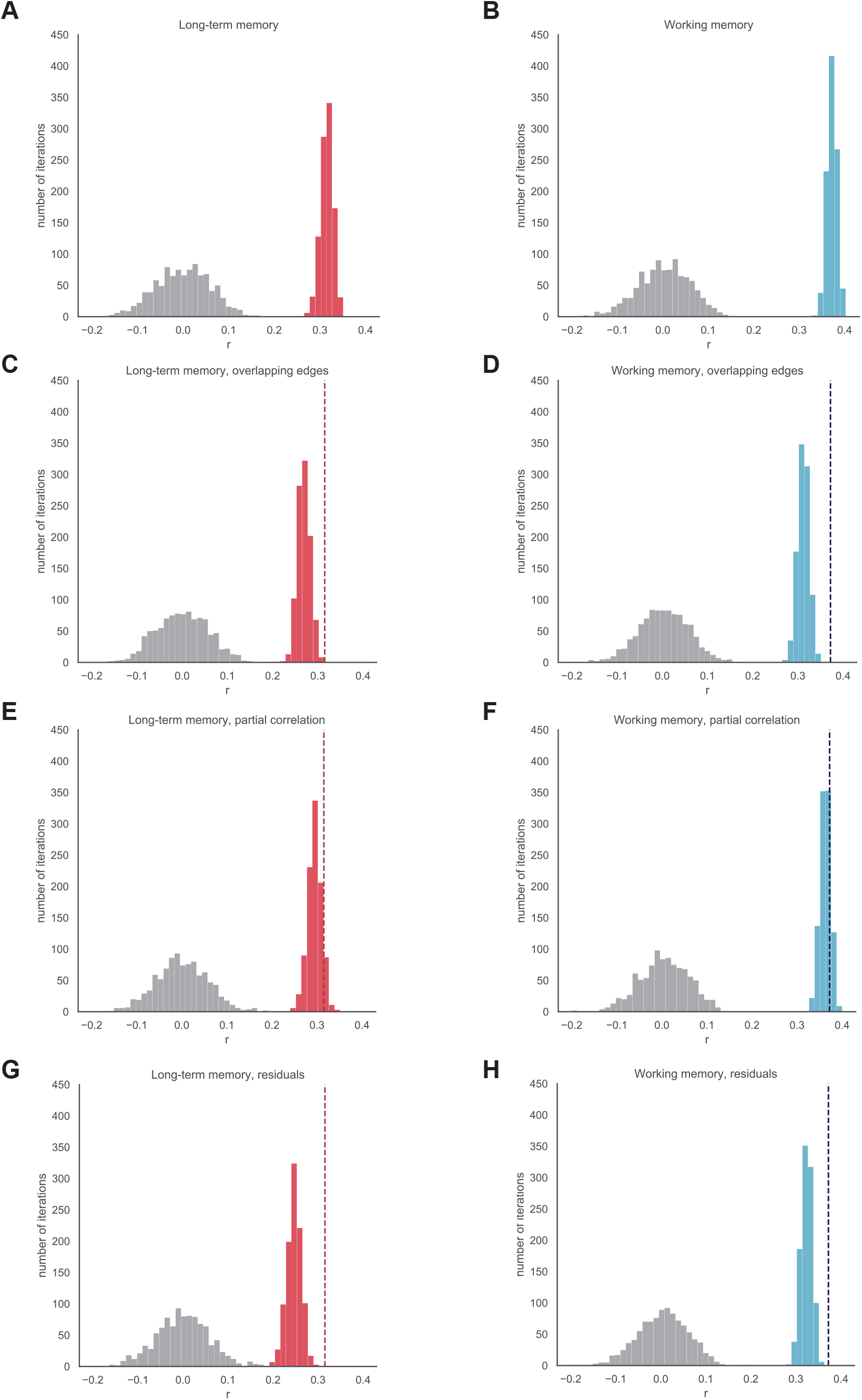
Distributions of true model performance (Red: LTM; Blue: WM) and null model performance (gray) for the different procedures reported in the main text. Model performance was measured by mean *r* between predicted and observed behavior across the 10 folds of each iteration with different partitions. Dotted lines represent the mean performance of the corresponding no-control models.

**Figure S5:**
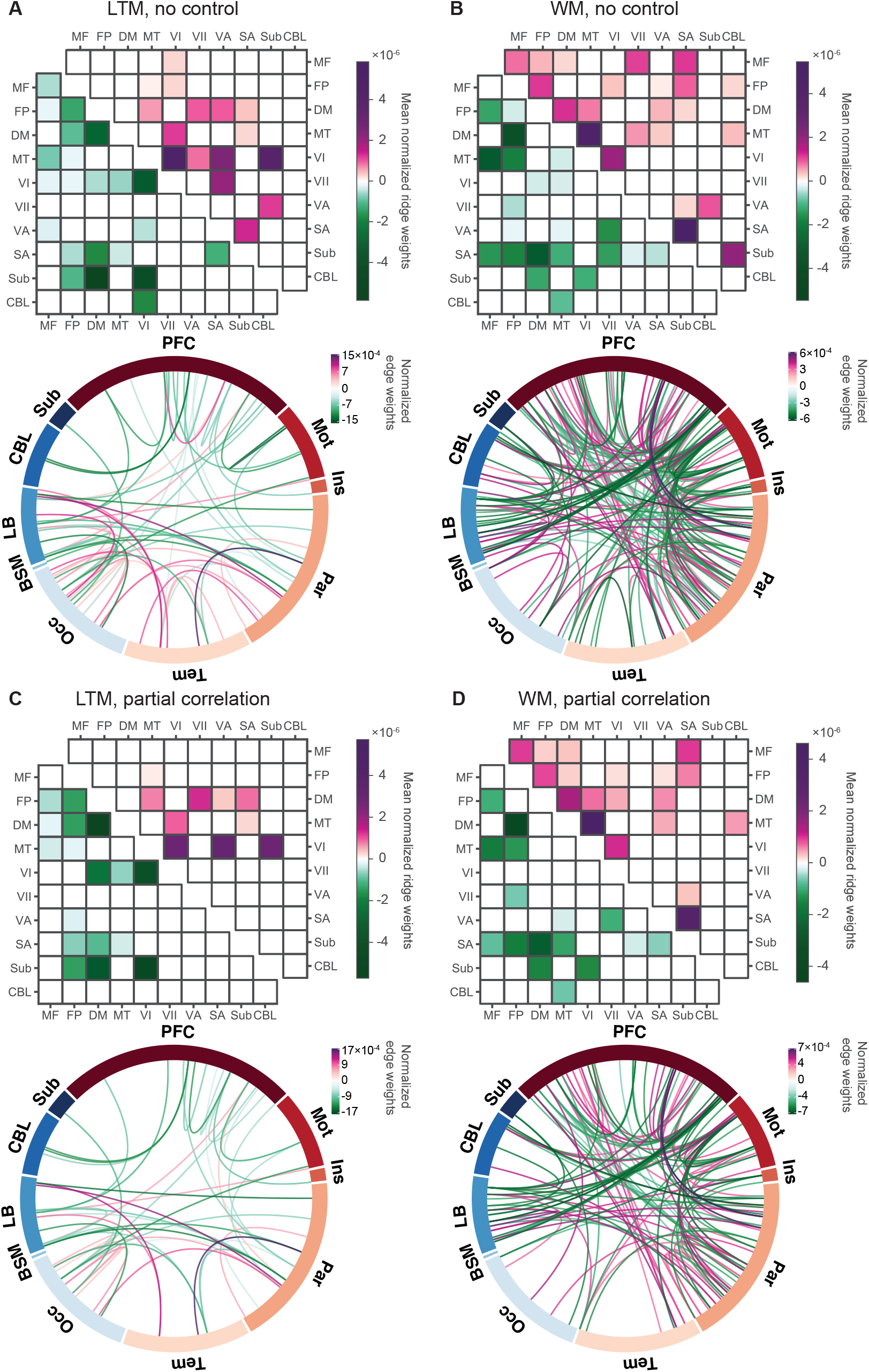
Anatomical distributions of predictive features for the no control models and the partial correlation models. For each panel, on the top are results from grouping nodes at the network level. Purple: features with positive weights, green: features with negative weights, shade: relative model contribution. Network labels: MF - medial frontal, FP - frontoparietal, DM - default mode, MT - motor, VI - visual I, VII - visual II, VA - visual association, SA - salience, Sub - subcortical, CBL - cerebellum. On the bottom are results from grouping nodes in to macroscale brain regions. The entire circle represents all 368 nodes, divided into the macroscale brain regions. The different locations on each ribbon represent the different nodes that belong to this region. Each line represents a reliable predictive edge. Purple: features with positive weights, green: features with negative weights, shade: relative edge weight. Macroscale regions labels: PFC - prefrontal cortex, Mot - motor cortex, Ins - insula, Par - parietal cortex, Tem - temporal cortex, Occ - occipital cortex, BSM - brainstem, LB – limbic regions (including the cingulate cortex, amygdala and hippocampus), Sub – subcortical regions (including the thalamus and striatum), CBL - cerebellum. (A) Predictive features of the model trained for LTM without controlling for WM. (B) Predictive features of the model trained for WM without controlling for LTM. (C) Predictive features of the model trained for LTM while controlling for WM during edge selection with partial correlation. (D) Predictive features of the model trained for WM while controlling for LTM during edge selection with partial correlation.

**Figure S6:**
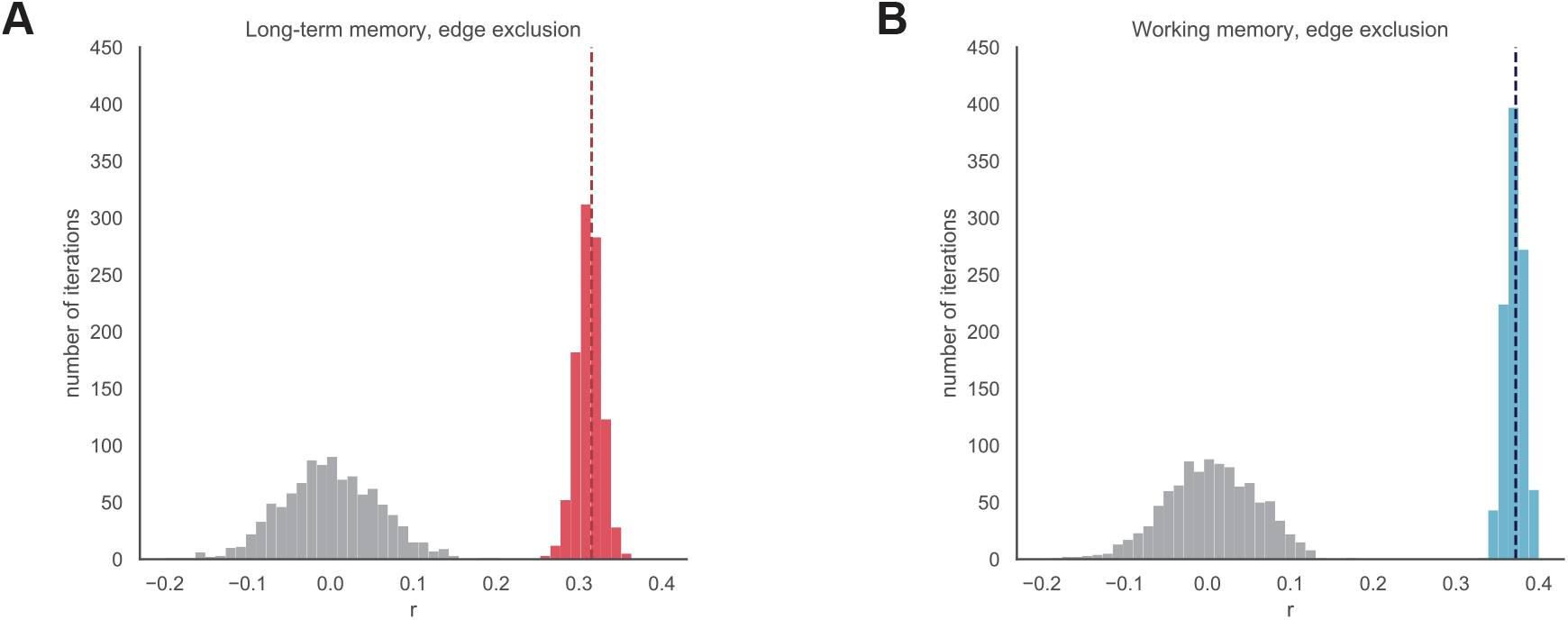
Model performance for predicting LTM and WM after controlling for the non-target behavior with edge exclusion (see methods for more details).

## Footnotes

1 We also ran the same analyses using resting state data but the LTM model performance was not significantly above chance (see Fig. S1) so we chose to focus on models built from n-back data here.

2 We did adopt another procedure to control for the non-target behavior during edge selection by explicitly excluding any edges that correlated with the target behavior (see Methods for details) and the prediction performance was very similar (see Fig. S6).

## References

Abraham, A., Pedregosa, F., Eickenberg, M., Gervais, P., Mueller, A., Kossaifi, J., Gramfort, A., Thirion, B., & Varoquaux, G. (2014). Machine learning for neuroimaging with scikit-learn. Frontiers in Neuroinformatics, 8, 14. https://doi.org/10.3389/fninf.2014.00014

Avery, E. W., Yoo, K., Rosenberg, M. D., Greene, A. S., Gao, S., Na, D. L., Scheinost, D., Constable, T. R., & Chun, M. M. (2019). Distributed Patterns of Functional Connectivity Predict Working Memory Performance in Novel Healthy and Memory-impaired Individuals. Journal of Cognitive Neuroscience, 1–15. https://doi.org/10.1162/jocn\_a\_01487

Barch, D. M., Burgess, G. C., Harms, M. P., Petersen, S. E., Schlaggar, B. L., Corbetta, M., Glasser, M. F., Curtiss, S., Dixit, S., Feldt, C., Nolan, D., Bryant, E., Hartley, T., Footer, O., Bjork, J. M., Poldrack, R., Smith, S., Johansen-Berg, H., Snyder, A. Z., … Consortium, W.-M. H. (2013). Function in the human connectome: Task-fMRI and individual differences in behavior. NeuroImage, 80, 169–189. https://doi.org/10.1016/j.neuroimage.2013.05.033

Bassett, D. S., & Sporns, O. (2017). Network neuroscience. Nature Neuroscience, 20 (3), 353–364. https://doi.org/10.1038/nn.4502

Bassett, D. S., Yang, M., Wymbs, N. F., & Grafton, S. T. (2015). Learning-induced autonomy of sensorimotor systems. Nature neuroscience, 18 (5), 744–751.

Bertolero, M. A., Yeo, B. T. T., Bassett, D. S., & D’Esposito, M. (2018). A mechanistic model of connector hubs, modularity and cognition. Nature Human Behaviour, 2 (10), 1803.08109, 765–777. https://doi.org/10.1038/s41562-018-0420-6

Blumenfeld, R. S., & Ranganath, C. (2006). Dorsolateral prefrontal cortex promotes long-term memory formation through its role in working memory organization. Journal of Neuroscience, 26 (3), 916–925.

Bullmore, E., & Sporns, O. (2009). Complex brain networks: graph theoretical analysis of structural and functional systems. Nature Reviews Neuroscience, 10 (3), 186–198. https://doi.org/10.1038/nrn2575

Conway, A. R. A., Kane, M. J., & Engle, R. W. (2003). Working memory capacity and its relation to general intelligence. Trends in Cognitive Sciences, 7 (12), 547–552.

Cowan, N. (2008). Progress in Brain Research. Progress in brain research, 169, 323–338. https://doi.org/10.1016/s0079-6123(07)00020-9

Dubois, J., & Adolphs, R. (2016). Building a Science of Individual Differences from fMRI. Trends in Cognitive Sciences, 20 (6), 425–443. https://doi.org/10.1016/j.tics.2016.03.014

Eichenbaum, H., Yonelinas, A. P., & Ranganath, C. (2007). The Medial Temporal Lobe and Recognition Memory. Annual Review of Neuroscience, 30 (1), 123–152. https://doi.org/10.1146/annurev.neuro.30.051606.094328

Essen, D. C. V., Smith, S. M., Barch, D. M., Behrens, T. E., Yacoub, E., Ugurbil, K., & Consortium, f. t. W.-M. H. (2013). The WU-Minn Human Connectome Project: An overview. NeuroImage, 80, 62–79. https://doi.org/10.1016/j.neuroimage.2013.05.041

Finn, E. S., Shen, X., Scheinost, D., Rosenberg, M. D., Huang, J., Chun, M. M., Papademetris, X., & Constable, R. T. (2015). Functional connectome fingerprinting: identifying individuals using patterns of brain connectivity. Nature Publishing Group, 18 (11), 1664–1671. https://doi.org/10.1038/nn.4135

Fischl, B. (2012). FreeSurfer. NeuroImage, 62 (2), 774–81. https://doi.org/10.1016/j.neuroimage.2012.01.021

Forbes, N. F., Carrick, L. A., McIntosh, A. M., & Lawrie, S. M. (2008). Working memory in schizophrenia: a meta-analysis. Psychological Medicine, 39 (6), 889–905. https://doi.org/10.1017/s0033291708004558

Gabrieli, J. D. E. (1996). Memory systems analyses of mnemonic disorders in aging and age-related diseases. Proceedings of the National Academy of Sciences, 93 (24), 13534–13540. https://doi.org/10.1073/pnas.93.24.13534

Gabrieli, J. D., Ghosh, S. S., & Whitfield-Gabrieli, S. (2015). Prediction as a humanitarian and pragmatic contribution from human cognitive neuroscience. Neuron, 85 (1), 11–26.

Gao, S., Greene, A. S., Constable, R. T., & Scheinost, D. (2019). Combining multiple connectomes improves predictive modeling of phenotypic measures. NeuroImage, 201, 116038. https://doi.org/10.1016/j.neuroimage.2019.116038

Glasser, M. F., Sotiropoulos, S. N., Wilson, J. A., Coalson, T. S., Fischl, B., Andersson, J. L., Xu, J., Jbabdi, S., Webster, M., Polimeni, J. R., Essen, D. C. V., Jenkinson, M., & Consortium, f. t. W.-M. H. (2013). The minimal preprocessing pipelines for the Human Connectome Project. NeuroImage, 80, 105–124. https://doi.org/10.1016/j.neuroimage.2013.04.127

Graham, K. S., Barense, M. D., & Lee, A. C. H. (2010). Going beyond LTM in the MTL: A synthesis of neuropsychological and neuroimaging findings on the role of the medial temporal lobe in memory and perception. Neuropsychologia, 48 (4), 831–853. https://doi.org/10.1016/j.neuropsychologia.2010.01.001

Greene, A. S., Gao, S., Noble, S., Scheinost, D., & Constable, R. T. (2020). How Tasks Change Whole-Brain Functional Organization to Reveal Brain-Phenotype Relationships. Cell Reports, 32 (8), 108066. https://doi.org/10.1016/j.celrep.2020.108066

Hsu, W.-T., Rosenberg, M. D., Scheinost, D., Constable, R. T., & Chun, M. M. (2018). Resting-state functional connectivity predicts neuroticism and extraversion in novel individuals. Social Cognitive and Affective Neuroscience, 13 (2), 224–232. https://doi.org/10.1093/scan/nsy002

James, W. (1891). The Principles of Psychology. The American Journal of Psychology, 3 (4), 578. https://doi.org/10.2307/1412102

Jangraw, D. C., Gonzalez-Castillo, J., Handwerker, D. A., Ghane, M., Rosenberg, M. D., Panwar, P., & Bandettini, P. A. (2018). A functional connectivity-based neuromarker of sustained attention generalizes to predict recall in a reading task. NeuroImage, 166 (Proc. Natl. Acad. Sci. U. S. A. 109 9 2012), 9–109. https://doi.org/10.1016/j.neuroimage.2017.10.019

Jeneson, A., & Squire, L. R. (2012). Working memory, long-term memory, and medial temporal lobe function. Learning & Memory, 19 (1), 15–25. https://doi.org/10.1101/lm.024018.111

Jenkinson, M., Beckmann, C. F., Behrens, T. E. J., Woolrich, M. W., & Smith, S. M. (2011). FSL. NeuroImage, 62 (2), 782–90. https://doi.org/10.1016/j.neuroimage.2011.09.015

Keerativittayayut, R., Aoki, R., Sarabi, M. T., Jimura, K., & Nakahara, K. (2018). Large-scale network integration in the human brain tracks temporal fluctuations in memory encoding performance. eLife, 7, e32696. https://doi.org/10.7554/elife.32696

Kim, H. (2011). Neural activity that predicts subsequent memory and forgetting: A meta-analysis of 74 fMRI studies. NeuroImage, 54 (3), 2446–2461. https://doi.org/10.1016/j.neuroimage.2010.09.045

Lever, A. G., Werkle-Bergner, M., Brandmaier, A. M., Ridderinkhof, K. R., & Geurts, H. M. (2015). Atypical working memory decline across the adult lifespan in autism spectrum disorder? Journal of Abnormal Psychology, 124 (4), 1014–1026. https://doi.org/10.1037/abn0000108

Lin, Q., Rosenberg, M. D., Yoo, K., Hsu, T. W., O’Connell, T. P., & Chun, M. M. (2018). Resting-State Functional Connectivity Predicts Cognitive Impairment Related to Alzheimer’s Disease. Frontiers in Aging Neuroscience, 10, 94. https://doi.org/10.3389/fnagi.2018.00094

Luck, S. J., & Vogel, E. K. (2013). Visual working memory capacity: from psychophysics and neurobiology to individual differences. Trends in Cognitive Sciences, 17 (8), 391–400. https://doi.org/10.1016/j.tics.2013.06.006

McNab, F., & Klingberg, T. (2008). Prefrontal cortex and basal ganglia control access to working memory. Nature Neuroscience, 11 (1), 103–107. https://doi.org/10.1038/nn2024

Meskaldji, D.-E., Preti, M. G., Bolton, T. A., Montandon, M.-L., Rodriguez, C., Morgenthaler, S., Gian-nakopoulos, P., Haller, S., & Ville, D. V. D. (2016). Prediction of long-term memory scores in MCI based on resting-state fMRI. NeuroImage: Clinical, 12 (J. Clin. Exp. Neuropsychol. 29 5 2007), 785795. https://doi.org/10.1016/j.nicl.2016.10.004

Noble, S., Spann, M. N., Tokoglu, F., Shen, X., Constable, R. T., & Scheinost, D. (2017). Influences on the test–retest reliability of functional connectivity mri and its relationship with behavioral utility. Cerebral Cortex, 27 (11), 5415–5429.

Olson, I. R., Moore, K. S., Stark, M., & Chatterjee, A. (2006a). Visual Working Memory Is Impaired when the Medial Temporal Lobe Is Damaged. Journal of Cognitive Neuroscience, 18 (7), 1087–1097. https://doi.org/10.1162/jocn.2006.18.7.1087

Olson, I. R., Page, K., Moore, K. S., Chatterjee, A., & Verfaellie, M. (2006b). Working Memory for Conjunctions Relies on the Medial Temporal Lobe. Journal of Neuroscience, 26 (17), 4596–4601. https://doi.org/10.1523/jneurosci.1923-05.2006

Paller, K. A., & Wagner, A. D. (2002). Observing the transformation of experience into memory. Trends in Cognitive Sciences, 6 (2), 93–102. https://doi.org/10.1016/s1364-6613(00)01845-3

Park, D. C., Smith, A. D., Lautenschlager, G., Earles, J. L., Frieske, D., Zwahr, M., & Gaines, C. L. (1996). Mediators of long-term memory performance across the life span. Psychology and Aging, 11 (4), 621–637. https://doi.org/10.1037//0882-7974.11.4.621

Park, D. C., Lautenschlager, G., Hedden, T., Davidson, N. S., Smith, A. D., & Smith, P. K. (2002). Models of visuospatial and verbal memory across the adult life span. Psychology and Aging, 17 (2), 299–320. https://doi.org/10.1037/0882-7974.17.2.299

Petersen, S. E., & Sporns, O. (2015). Brain networks and cognitive architectures. Neuron, 88 (1), 207–219.

Power, J. D., Mitra, A., Laumann, T. O., Snyder, A. Z., Schlaggar, B. L., & Petersen, S. E. (2014). Methods to detect, characterize, and remove motion artifact in resting state fmri. Neuroimage, 84, 320–341.

Ranganath, C., & Blumenfeld, R. S. (2005). Doubts about double dissociations between short-and long-term memory. Trends in Cognitive Sciences, 9 (8), 374–380. https://doi.org/10.1016/j.tics.2005.06.009

Ranganath, C., Cohen, M. X., & Brozinsky, C. J. (2005). Working Memory Maintenance Contributes to Long-term Memory Formation: Neural and Behavioral Evidence. Journal of Cognitive Neuroscience, 17 (7). https://doi.org/10.1162/0898929054475118

Ranganath, C., & D’Esposito, M. (2001). Medial temporal lobe activity associated with active maintenance of novel information. Neuron, 31 (5), 865–873.

Rosenberg, M. D., Finn, E. S., Scheinost, D., Papademetris, X., Shen, X., Constable, R. T., & Chun, M. M. (2015). A neuromarker of sustained attention from whole-brain functional connectivity. Nature Neuroscience, 19 (1), 165–171. https://doi.org/10.1038/nn.4179

Rosenberg, M. D., Scheinost, D., Greene, A. S., Avery, E. W., Kwon, Y. H., Finn, E. S., Ramani, R., Qiu, M., Constable, R. T., & Chun, M. M. (2020). Functional connectivity predicts changes in attention observed across minutes, days, and months. Proceedings of the National Academy of Sciences, 117 (7), 3797–3807.

Rugg, M. D., & Vilberg, K. L. (2013). Brain networks underlying episodic memory retrieval. Current Opinion in Neurobiology, 23 (2), 255–260. https://doi.org/10.1016/j.conb.2012.11.005

Scheinost, D., Noble, S., Horien, C., Greene, A. S., Lake, E. M., Salehi, M., Gao, S., Shen, X., O’Connor, D., Barron, D. S., Yip, S. W., Rosenberg, M. D., & Constable, R. T. (2019). Ten simple rules for predictive modeling of individual differences in neuroimaging. NeuroImage, 193 (Neuroimage 147 2017), 35–45. https://doi.org/10.1016/j.neuroimage.2019.02.057

Scoville, W. B., & Milner, B. (1957). Loss of recent memory after bilateral hippocampal lesions. Journal of neurology, neurosurgery, and psychiatry, 20 (1), 11.

Shen, X., Tokoglu, F., Papademetris, X., & Constable, R. T. (2013). Groupwise whole-brain parcellation from resting-state fMRI data for network node identification. NeuroImage, 82, 403–415. https://doi.org/10.1016/j.neuroimage.2013.05.081

Shen, X., Finn, E. S., Scheinost, D., Rosenberg, M. D., Chun, M. M., Papademetris, X., & Constable, R. T. (2017). Using connectome-based predictive modeling to predict individual behavior from brain connectivity. Nature Protocols, 12 (3), 506–518. https://doi.org/10.1038/nprot.2016.178

Shine, J. M., Breakspear, M., Bell, P. T., Martens, K. A. E., Shine, R., Koyejo, O., Sporns, O., & Poldrack, R. A. (2019). Human cognition involves the dynamic integration of neural activity and neuromodulatory systems. Nature neuroscience, 22 (2), 289–296.

Snyder, H. R. (2013). Major depressive disorder is associated with broad impairments on neuropsychological measures of executive function: A meta-analysis and review. Psychological Bulletin, 139 (1), 81–132. https://doi.org/10.1037/a0028727

Squire, L. R., Stark, C. E. L., & Clark, R. E. (2004). THE MEDIAL TEMPORAL LOBE. Annual Review of Neuroscience, 27 (1), 279–306. https://doi.org/10.1146/annurev.neuro.27.070203.144130

Süß, H.-M., Oberauer, K., Wittmann, W. W., Wilhelm, O., & Schulze, R. (2002). Working-memory capacity explains reasoning ability—and a little bit more. Intelligence, 30 (3), 261–288. https://doi.org/10.1016/s0160-2896(01)00100-3

Tambini, A., & Davachi, L. (2019). Awake Reactivation of Prior Experiences Consolidates Memories and Biases Cognition. Trends in Cognitive Sciences. https://doi.org/10.1016/j.tics.2019.07.008

Tambini, A., Ketz, N., & Davachi, L. (2010). Enhanced brain correlations during rest are related to memory for recent experiences. Neuron, 65 (2), 280–290. https://doi.org/10.1016/j.neuron.2010.01.001

Todd, J. J., & Marois, R. (2004). Capacity limit of visual short-term memory in human posterior parietal cortex. Nature, 428 (6984), 751–754. https://doi.org/10.1038/nature02466

Todd, J. J., & Marois, R. (2005). Posterior parietal cortex activity predicts individual differences in visual short-term memory capacity. Cognitive, Affective, & Behavioral Neuroscience, 5 (2), 144–155. https://doi.org/10.3758/cabn.5.2.144

Trelle, A. N., Carr, V. A., Guerin, S. A., Thieu, M. K., Jayakumar, M., Guo, W., Nadiadwala, A., Corso, N. K., Hunt, M. P., Litovsky, C. P., et al. (2020). Hippocampal and cortical mechanisms at retrieval explain variability in episodic remembering in older adults. Elife, 9, e55335.

Turk-Browne, N. B. (2013). Functional Interactions as Big Data in the Human Brain, 1 6.

Unsworth, N. (2010). On the division of working memory and long-term memory and their relation to intelligence: A latent variable approach. Acta Psychologica, 134 (1), 16–28. https://doi.org/10.1016/j.actpsy.2009.11.010

Unsworth, N. (2019). Individual differences in long-term memory. Psychological Bulletin, 145 (1), 79.

Wagner, A. D., Schacter, D. L., Rotte, M., Koutstaal, W., Maril, A., Dale, A. M., Rosen, B. R., & Buckner, R. L. (1998). Building Memories: Remembering and Forgetting of Verbal Experiences as Predicted by Brain Activity. Science, 281 (5380), 1188–1191. https://doi.org/10.1126/science.281.5380.1188

Xu, Y., & Chun, M. M. (2006). Dissociable neural mechanisms supporting visual short-term memory for objects. Nature, 440 (7080), 91–95. https://doi.org/10.1038/nature04262

Yamashita, M., Yoshihara, Y., Hashimoto, R., Yahata, N., Ichikawa, N., Sakai, Y., Yamada, T., Matsukawa, N., Okada, G., Tanaka, S. C., Kasai, K., Kato, N., Okamoto, Y., Seymour, B., Takahashi, H., Kawato, M., & Imamizu, H. (2018). A prediction model of working memory across health and psychiatric disease using whole-brain functional connectivity. elifesciences.org, 7. https://doi.org/10.7554/elife.38844

Yarkoni, T., Poldrack, R. A., Nichols, T. E., Essen, D. C. V., & Wager, T. D. (2011). Large-scale automated synthesis of human functional neuroimaging data. Nature Methods, 8 (8), 665–670. https://doi.org/10.1038/nmeth.1635

